# In-section Click-iT detection and super-resolution CLEM: Shedding light on nucleolar ultrastructure and S-phase progression in plants

**DOI:** 10.1101/2023.05.10.540144

**Authors:** M Franek, L Koptasikova, J Miksatko, J Pospisil, D Liebl, M Esner, E Macickova, M Dvorackova, J Fajkus

## Abstract

Correlative light and electron microscopy (CLEM) is an essential tool that allows for localisation of a particular target molecule(s) and their spatial correlation with the ultrastructural map of subcellular features at the nanometer scale. Adoption of these advanced imaging methods has been limited in plant biology, due to challenges with plant tissue permeability, fluorescence labelling efficiency, indexing of features of interest throughout the complex 3D volume and their re-localization on micrographs of ultrathin cross-sections. Here, we demonstrate an imaging approach based on tissue processing and embedding into methacrylate resin followed by imaging of serial sections by both, single-molecule localization microscopy and transmission electron microscopy for correlative analysis. Importantly, we demonstrate that the use of a particular type of embedding resin is not only compatible with single-molecule localization microscopy but shows a dramatic improvement in fluorophore blinking behavior relative to the whole-mount approaches. Here we used commercially available Click-iT ethynyl-deoxyuridine cell proliferation kit to visualize the DNA replication sites of wild-type *Arabidopsis thaliana* seedlings, as well as FASCIATA1 and NUCLEOLIN1 mutants and applied our on-section CLEM imaging workflow for the analysis of S-phase progression and nucleolar organization in mutant plants with aberrant nucleolar phenotypes.

## Introduction

Correlative light and electron microscopy (CLEM) combines the unique benefits of light and electron microscopy (EM), where images of specific, fluorescently labelled cellular structures captured in fluorescence microscope are spatially correlated with a detailed map of ultrastructural features revealed by electron microscopy. In practice, correlative microscopy approaches often entail a number of challenges, namely in the choice of an appropriate chemical fixation, processing, resin embedding and labelling steps that have to suit both, preservation of fluorescence (or antigens for immunofluorescence staining) and preservation of fine ultrastructure for electron microscopy. Conventional chemical fixation, post-contrasting with osmium and dehydration prior to sample embedding in resin for transmission EM often results in partial extraction or denaturation of proteins, loss of fluorescence (in the case of fluorescent proteins) or leads to an inefficient post-embedding immuno-labelling due to the destruction or limited accessibility of epitopes on the surface of resin sections. The last decade brought forward advances in sample preparation for CLEM, which demonstrated the feasibility of ultrastructural analysis by EM while maintaining the fluorescence of endogenous proteins with the possibility of super-resolution imaging (Kopek et al., 2012, Johnson et al., 2015, Kim et al., 2015) or protocols that enable post-embedding on-section fluorescence labelling (Wurm et al., 2019). The combination of super-resolution microscopy (SRM), especially single-molecule localization microscopy (SMLM), with the EM brings about further technical considerations. SMLM uses the stochastic photo-switching (“blinking”) of fluorophores to determine their positions with resolution below the diffraction limit (∼5 – 40 nm), therefore retention of the blinking potential of fluorophores upon embedding in a resin as well as epitope accessibility and reactivity in resin sections are crucial. To date, few groups have reported successful implementations of SRM-CLEM (Watanabe et al., 2010, Hoffman et al., 2020), mostly on adherent mammalian cell lines (Kim et al., 2015; Johnson et al., 2015; Tujitel et al., 2019).

Implementation of these techniques in plants is not trivial, since there is no equivalent of mammalian cell culture systems and plant cell walls are a considerable obstacle for the highly-specific labelling of structures required for localization microscopy (Sauer et al., 2006; Pasternak et al., 2015). We sought to implement a sample preparation and imaging protocol for correlative microscopy that would retain the possibility to identify particular type of plant tissue, allow inspection of the ultrastructural features by the EM using common contrasting techniques, and simultaneously permit super-resolution imaging of DNA replication sites.

Novel methodological approaches in super-resolution microscopy/nanoscopy are generally benchmarked on well-described cellular structures, such as the nuclear pore complex or cytoskeletal components (Huang et al., 2008; Thevathasan et al., 2019). Here, we developed a correlative workflow to study nucleolar architecture changes during progression of DNA replication in plant cells. We have previously characterized the organization and replication of ribosomal DNA in plant nuclei (Dvorackova et al., 2018, Kutashev et al., 2020) using fluorescence in-situ hybridization and replication labelling. We have shown that the replication of the 45S ribosomal DNA (rDNA) occurs throughout S-phase in the nucleus and nucleolus, whereby the active intranucleolar fraction of rDNA replicates in early-S-phase, larger intranucleolar clusters of rDNA genes replicate in the mid S-phase and the perinucleolar, transcriptionally inactive fraction replicates in the late S-phase. Results from replication labelling combined with sequencing (Repli-SEQ) have shown that nucleolus displays a bipartite replication pattern, with distinct segments of rDNA replicating in the early and in the late S-phase, in agreement with microscopic observations (Concia et al., 2018, Wear et al., 2017).

To investigate the links between nucleolar architecture and replication progression in the nucleolus, we selected specific *A. thaliana* mutants, known for alterations in nucleolar structure (*nuc1*) or progressive loss of ribosomal genes (*fas*1). It has been previously reported that *nuc*1 mutants display changes in nucleolar architecture (Pontvianne et al., 2007; Pontvianne et al., 2010, Durut et al., 2014), with changes in the methylation pattern of rDNA genes. *Fas*1 mutants have been shown to have a decreased number of rDNA copies and exhibit general chromatin decompaction (Muchova et al., 2015; Kolarova et al., 2020).

In this work, we establish a novel correlative imaging approach in tissue sections of the *A. thaliana*, integrating the sequential analysis of replication with SMLM and of nucleolar structure by transmission electron microscopy (TEM), with the localization precision in the range of 10-20 nm for SMLM data. We demonstrate that (i) imaging of plant tissue segments embedded in Lowicryl is feasible by both, SMLM and TEM; (ii) SMLM performed on this type of sections has high quality metrics and (iii) these sections are permeable for low molecular weight (MW) labelling compounds (Click-iT chemistry, phalloidin labelling) and allows for labelling throughout the section volume. We demonstrate that rDNA replication patterns in plants can be precisely localised and visualized by (pre-)labelling of samples with ethynyl-5 ‘-deoxyuridine (EdU) followed chemical processing, resin embedding and performing Click-iT chemistry detection directly on semithin sections. Moreover, serial sectioning of alternating semithin (500 nm) and ultrathin sections (70 nm) allows us to probe the ultrastructure of the nucleolus with TEM and the localization of fluorescently labelled replication patterns in the adjacent cell section in parallel, with a negligible offset in the z-axis. Our data suggest that the presence of intranucleolar replication foci (fluorescent labelling) is concomitant with the detection of fibrillar centers in the nucleoli (discernible on TEM micrographs) and the differences in the nucleolar architecture correlate with changes in the size of intranucleolar replication foci between wt, *fas1* and *nuc1* mutants.

## Materials and Methods

### Plant growth and EdU incubation

*Arabidopsis thaliana* seeds were sterilized (90% ethanol, 5 min) and plated on half-strength agar Murashige and Skoog medium (½ MS medium) with 1% sucrose. After 1-day stratification (4°C/dark) plates were transferred to the growth chamber and grown for up to 1 week under long day (LD) conditions (16 h light - 21°C/8 h dark - 19°C/50-60% relative humidity). 7-day-old seedlings of wt Col0, *fas1*, and *nuc1* mutants were incubated in 20uM EdU in 1x liquid MS medium for 90 min and subsequently processed for light and electron microscopy as described below.

### Progressive lowering of temperature and embedding in Lowicryl resin

After the EdU incubation, the samples were rinsed in 1x PHEM buffer (60mM PIPES, 25mM HEPES, 10mM EGTA, 2mM MgCl2) pH 6.8 and fixed in 4% formaldehyde (FA, #15710, EMS, Hatfield, USA) and 0.1% glutaraldehyde (GA, #16220, EMS, Hatfield, USA) in PHEM buffer on a rotating platform for 1 hour at room temperature, then overnight at 4°C. After three washes with cold PHEM buffer pH=6.8 for 10 min, the samples were embedded into 2% low-melting agarose (LMA; 2% agarose in ddH_2_O) blocks (according to Wu et al., 2012) and *en block* contrasted with 0.5% uranyl acetate aqueous solution at 4°C overnight. Dehydration and lowicryl K4M (#14330, EMS, Hatfield, USA) infiltration were performed manually using progressive lowering of temperature technique in the freeze-substitution unit (Leica EM AFS2). Increasing concentration of ethanol was accompanied by decreasing temperature in the chamber down to -35°C. During this process, the ethanol was substituted with LowicrylK4M resin at different ratios, and finally infiltrated with 100% Lowicryl K4M. The samples were polymerized under UV at -35°C for three days.

### Microwave-assisted processing to Spurr’s resin

For the ultrastructural analysis, the samples were rinsed in 1x PHEM buffer and fixed in 3% GA and 0.5% FA in PHEM buffer, pH 6.8. After incubation for 3 hours at room temperature, the samples were fixed in 0.5 % GA and 4% FA in PHEM buffer pH 6.8 and stored at 4°C overnight until further processing. Before embedding to 2% LMA, the samples were washed in cold PHEM buffer at room temperature and all subsequent steps were conducted by a microwave tissue processor (Pelco BioWave Pro+ #3670-230, Redding, USA) equipped with power modulator, vacuum sample container and ColdSpot (system preventing local hotspots generation). All microwave steps (except for dehydration) were performed under vacuum. Samples were first contrasted with osmium tetroxide (0.5% in miliQ water) while microwaved at a power of 100W in 2 min ON/OFF cycles for 14 min. This was followed by three washes in PHEM buffer and contrasting in 0.5% uranyl acetate at a power of 150W for 7 min. Gradual dehydration in acetone was performed at a power of 250W for 1 min at each step, followed by infiltration in Spurr’s resin (#14300, EMS, Hatfield, USA) at ratios 1:3, 1:1, 3:1 with acetone and three changes of pure 100% Spurr’s resin at a power of 150W for 7 min each step. The specimens were finally polymerised at 70°C for 72 h in flat embedding silicon molds (#70905-01, EMS, Hatfield, USA).

### Collection of semithin and ultrathin sections

For correlative microscopy, cutting and collection of each semithin (500nm) section was followed by cutting and collection of an adjacent ultrathin (70nm) section to reduce the Z-offset in image correlation. The ultrathin sectioning was performed in the ultramicrotome (Leica EM UC7) using Ultra Sonic Diamond knife (angle 35°, Diatome, Hatfield, Switzerland). The semithin sections were collected onto glass coverslips coated with 0.05% poly-L-lysine (#P8920; Sigma-Aldrich), ultrathin sections on carbon-formvar-coated copper-slot-grids (#G2010-Cu, EMS, Hatfield, USA). Thicker sections (500nm and 1000nm) were collected to test resin permeability by labelling for 3D-SMLM. All semithin sections for super-resolution microscopy were labelled using Click-iT chemistry, ultrathin sections for TEM were post-contrasted using aqueous 4% uranyl acetate (#22400, EMS, Hatfield, USA) and 3% Reynold’s lead citrate.

### Click-iT labelling, phalloidin staining and immunolabelling

Click-iT detection of EdU on sections attached to poly-L-lysine coated coverslips was performed using the Click-iT cell proliferation kit (#C10340; Thermo Fisher Scientific), according to the manufacturer’s protocol. For the staining of actin filaments, sections on coverslips were first washed two times in 1x PBST (0.05% Tween-20), blocked for 30 minutes in 3% BSA in 1x PBST and incubated with 150 nM Phalloidin-AF647 (#A22287, Thermo Fisher Scientific) in 3% BSA / PBST for 30 minutes in a humid chamber. Samples were then rinsed 3x in 1x PBST and imaged in the imaging buffer consisting of 50 mM Tris-HCl, 10 mM NaCl, 50 mM β-mercaptoethylamine (MEA, 30070, Sigma-Aldrich), 1.1 mg/ml glucose oxidase (G2133, Sigma-Aldrich) and 100 μg/ml catalase (C40, Sigma-Aldrich, St. Louis, MO, USA) in Chamlide magnetic holder cells (#CM-B25-1; Live Cell Instrument). For immunolabelling of histone H3, sections on coverslips were first blocked for 30 minutes in 3% BSA in 1x PBST, then incubated with anti-H3 antibody (1:100 dilution, #ab1791, Abcam) in BSA / PBST overnight (#ab1791; Abcam) followed by rinse in 1x PBST and incubation with Alexa Fluor Plus 555 secondary antibody (1:200 dilution, #A32732, Thermo Fisher Scientific) for 45 minutes at room temperature prior to final wash in 1x PBST.

### Fluorescence light microscopy

The images of 500 nm thick root sections were acquired on the Nikon CSU-W1 confocal spinning disk microscope. Time-lapse dSTORM imaging was performed on a Nikon Ti-E microscope with a Nikon CFI HP Apo TIRF 100x oil objective (1.49 NA, detection with EM CCD Andor iXon Ultra DU897 camera) and on Zeiss Elyra 7 (Carl Zeiss, GmBH) imaging system with a 63x Oil Plan-Apochromat oil objective (1.46 NA, detection with PCO edge 4.2 sCMOS camera) in the HILO mode. For SMLM of whole-mount samples, roots were immobilized on poly-L-lysine coated high-precision coverslips and mounted in an imaging buffer (as described above). For SMLM of sections, 500 nm and 1000 nm thick root sections were anchored to 22×22 mm high-precision poly-L-lysine coated coverslips, mounted in Chamlide holder cells and imaged as described above.

### Electron microscopy

TEM images of ultrathin (70 nm) sections were acquired on the Jeol JEM-2100Plus (200kV) equipped with LaB_6_ cathode, TVIPS XF416 CMOS 4kx4k camera and SerialEM software v. 4.0.3 (Mastronarde, 2005). Montages of root tips (each consisting of several hundred images) were collected at pixel size of 2.83 nm, 0.7 s exposure, and 2x binning, in order to facilitate the localisation of target cells for cross-correlation with images from brightfield and fluorescence microscopy. Details were recorded with different pixel sizes depending on the region of interest, 2 s exposure and 1x binning. Data were aligned and pre-processed in IMOD software v. 4.11.4 (Kremer et al., 1996).

### Image analysis and SMLM reconstructions

For SMLM reconstructions, 20 000 time frames were acquired. Reconstructions of super-resolution images were performed using ThunderSTORM (Ovesny et al., 2016). To validate the quality of the SMLM reconstructions by quantitative error mapping of super-resolution images we used the NanoJ-SQUIRREL package (Culley et al., 2018; Schindelin et al., 2012) in FIJI. For Fourier ring correlation (FRC) analysis, 20 000 images from the time series were split into odd and even frames, which were used to reconstruct super-resolution images in ThunderSTORM. Images were then merged into a stack in FIJI and used as the input for the FRC analysis in NanoJ-SQUIRREL. Density-based cluster analysis was performed in the SMAP environment (Ries, 2020). For the clustering analysis, the super-resolution reconstructions were first processed by the density calculator (counting neighbours for each localization in a circle < 12 nm), which was then used to filter out localizations in lowest-density regions. Subsequently, we calculated the number of clusters using DBscan, using the SMAP implementation of the algorithm by Caetano et al., 2015; with the minimum number of objects in the neighbourhood set to 7; the neighbouring radius set to 20. 3D surface projections were rendered using the Imaris software (Bitplane, Oxford Instruments).

### Statistical analysis

For quantitative analysis, we pooled data from two biological replicates (e.g. two different seedlings/condition) and two technical replicates for each biological replicate (e.g. different sections contrasted/labelled on a coverslip or EM grid). For the statistical analysis of the size of intranucleolar replication foci (IRF) and fibrillar centres (FCs), we applied the non-parametric Kruskal-Wallis H-test and subsequently the Wilcoxon rank sum post-hoc test. H-values, p-values and adjusted p-values for post-hoc tests are presented in the text. Data normality was checked using the Shapiro-Wilk test for normality. Statistical analyses were performed in the Scipy python library.

## Results

### Correlative workflow for large plant tissue segments

To access the ultrastructure of the plant nucleolus in relation to the S-phase progression, we aimed to perform Click-iT chemistry labelling on semithin sections of plant roots embedded in EM resin. First, we incubated seedlings with EdU, fixed the samples by chemical cross-linking and embedded samples into low-melting agarose blocks for easier handling of the roots (stem and leaves trimmed off) during subsequent steps of chemical processing, as reported in Wu et al., 2012 (Fig. 1 A, B). Next, two types of resin were tested for sample embedding and *on-section* Click-iT labelling: (i) Spurr’s resin, an epoxy-based, high-penetration, low-viscosity, hydrophobic resin and (ii) Lowicryl K4M, a methacrylate-based, water-compatible, polar resin (referred to further in the text as “Lowicryl sections” for simplicity; Fig. 1 C, D). The Click-iT replication labelling was then performed on 500 nm longitudinal sections prepared from the resin-embedded seedlings, (Fig. 1E) while the uranyl acetate and lead citrate post-contrasting was performed on alternate 70 nm sections (Fig. 1F). The best signal-to-noise ratio was achieved in Lowicryl-embedded samples (Fig. 2A), with discernible differences in EdU patterns reflecting the different stages of DNA replication (Fig. 2E, e1-e3). Embedding of samples into Spurr’s resin, which includes conventional osmium tetroxide contrasting, is compatible with Click-iT labelling on resin sections (Fig. 2 C, D). In this case, the signal-to-noise ratio was much lower when compared with labelling of sections from Lowicryl embedded samples (Fig. 2 A, B).

**Figure 1.**
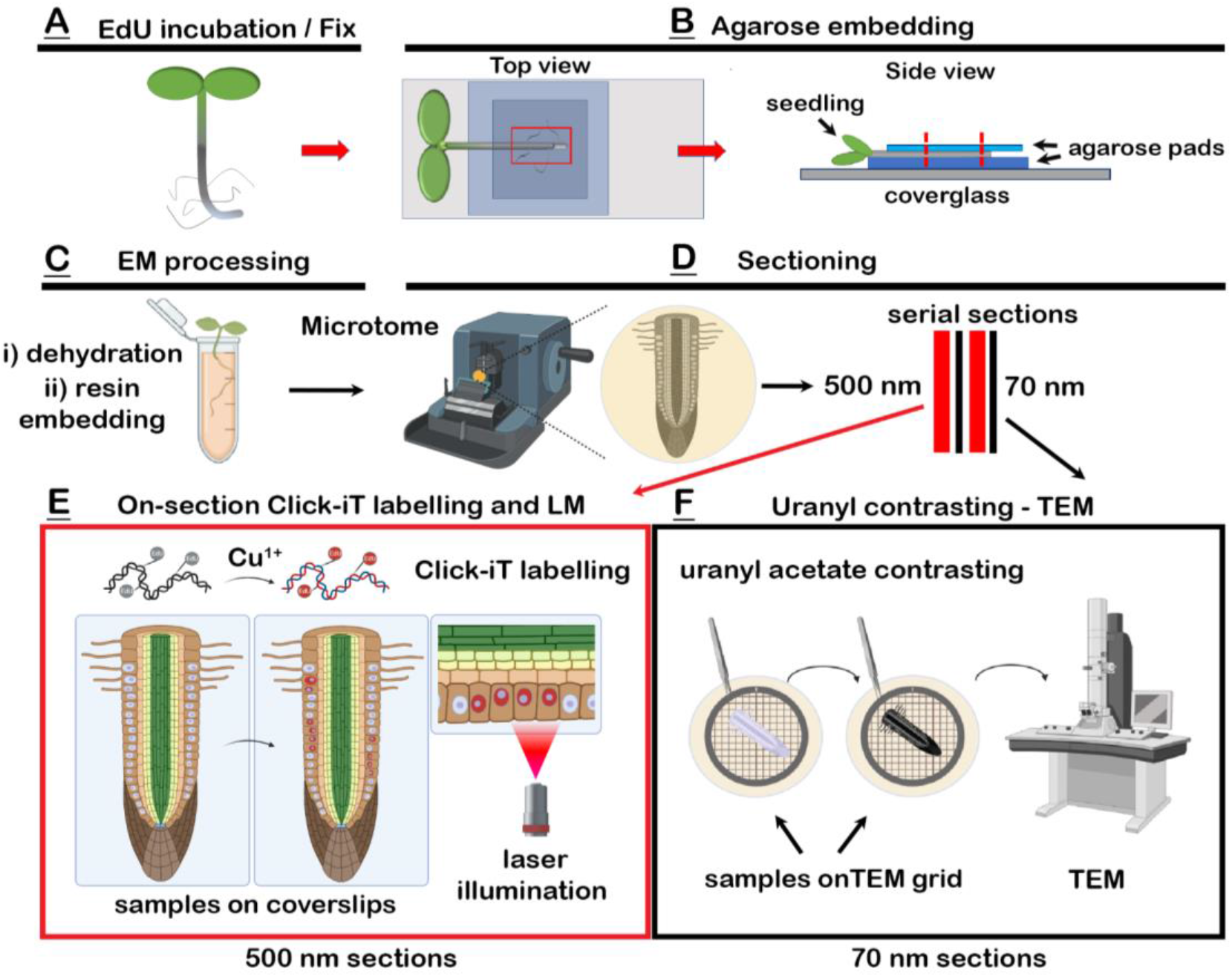
Schematic representation of the CLEM workflow for *A. thaliana*. **A**) Seedlings were labelled with EdU, fixed in a mixture of glutaraldehyde and formaldehyde and embedded into a block of low-melting agarose (**B**). Strips of agarose-embedded roots were post-fixed, dehydrated and embedded into Spurr’s or Lowicryl resin (**C**). Polymerized resin blocks were sectioned (**D**) with an ultramicrotome into consecutive pairs of semi-thin (500 nm) sections for fluorescence microscopy and ultra-thin sections (70 nm) for electron microscopy. Click-iT labelling was performed on 500 nm sections (**E**) and images were captured by either spinning disk or single-molecule localization microscopy. Ultrathin sections were collected on EM slots, post-contrasted with 4% uranyl acetate and imaged in a Transmission electron microscope (TEM) (**F**). Legend: **LM** - light microscopy, **TEM** - transmission electron microscopy.

**Figure 2.**
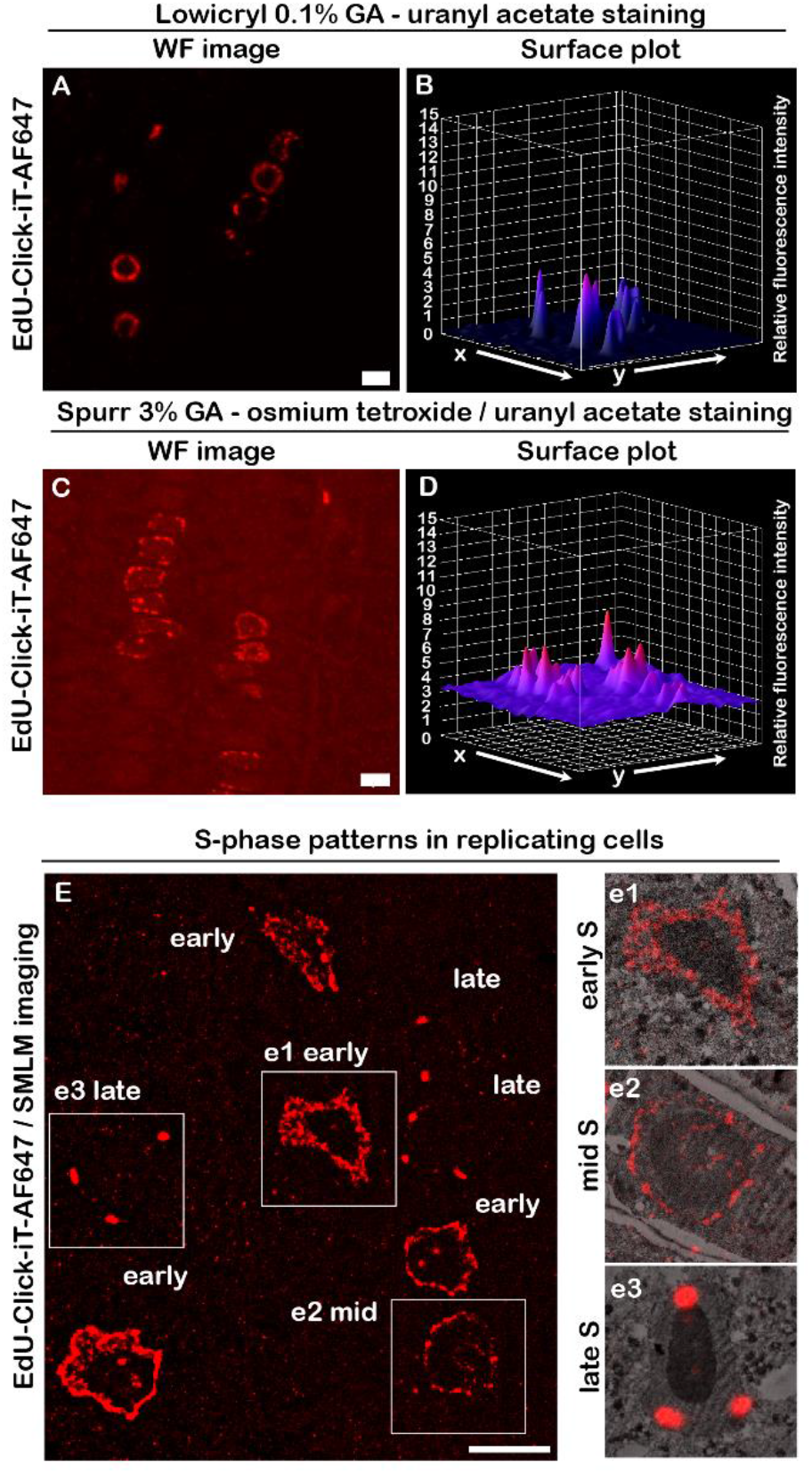
Replication labelling with EdU and Click-iT chemistry in Spurr and Lowicryl embedded samples. Plant root sections embedded in Lowicryl (**A**) and Spurr (**C**) resin labelled with Click-iT chemistry, with surface plots showing the relative background levels in the respective fixation and embedding conditions (**B, D**). **E**) Large field of view reconstruction of the SMLM image highlighting cells in different stages of DNA replication. Nuclei showing distinct replication patterns (**e1** – early-S-phase, **e2** – mid-S-phase and **e3** – late-S-phase) are displayed in an overlay with the corresponding TEM micrographs. Scale bar: A, C – 50 μm, E - 5 μm.

Having established that resin embedding does not preclude Click-iT detection, we aimed to analyse whether image registration is possible between fluorescence and EM micrographs capturing large segments of plant tissue sections (Fig. 3 A, B). In general, image registration for CLEM is accomplished either by mapping defined cellular structures (e.g. labelled mitochondria; Hoffman et al., 2020) or using fluorescently labelled gold nanoparticles as fiducial markers (Watanabe et al., 2010, Kopek et al., 2012). To align the images from electron microscopy and light microscopy, we took advantage of the anatomical features of the root and cell wall morphology, visible in both modalities (Fig. 3 C, D). Consistent with previously published literature (Hoffman et al., 2020; Kopek et al., 2012), we noted shrinking, stretching and distortion artifacts between sections (Fig. 3E, F). Ultimately, we were able to correlate replication patterns obtained from fluorescence microscopy with the corresponding TEM micrographs locally for individual cells (Fig. 3 G - I). Having demonstrated the possibility of Click-iT labelling on Lowicryl sections in a correlative workflow, we next sought to establish whether we could map the ultrastructural details of replicating chromatin segments by implementing SMLM in cells tagged with Click-iT chemistry.

**Figure 3.**
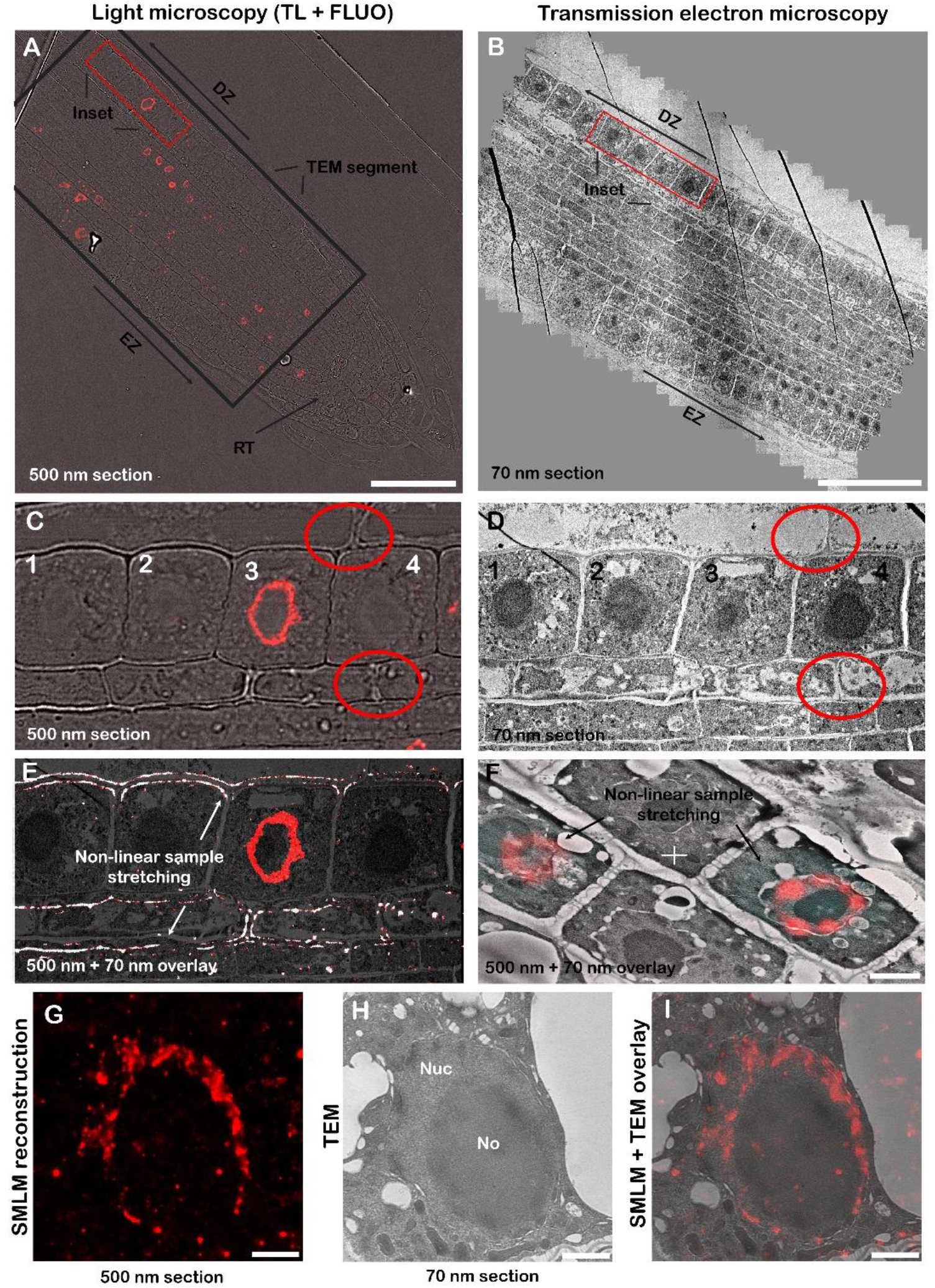
Correlative light-electron imaging of *A. thaliana* roots. Spinning-disk fluorescence (**A**) and transmission electron microscopy (**B**) imaging of roots on 500 nm and 70 nm sections, respectively. Composite (panorama) image of individual EM fields of view to capture an overview of a large area of the root (**B**). Insets, highlighted by red rectangles, magnified in **C** and **D**, showcasing cells of interest (**1**-**4**) in the differentiation zone of the root, with anatomical features of the root used for cell identification highlighted by red circles. Anisotropic sample stretching is shown in an overlay CLEM image (**E**, confocal image thresholded to highlight cell walls). Arrows highlight the problems with image registration, shown on cell wall architecture (notice the mismatch of cell wall contours at the bottom and good fitting at top of the image). **F)** Anisotropic stretching shown in the displacement of the fluorescence signal from nuclei, not fitting the underlying cell localization observed in TEM data. **G**) Detailed view of a root nucleus shown in SMLM reconstruction. Details of the same cell in the corresponding EM image (**H**) and image overlay shown in (**I**). Legend: **TL** - transmitted light, **FLUO** - fluorescence, **EZ** - elongation zone, **DZ** – differentiation zone, **RT** – root tip, **Nuc** - nucleus, **No** - nucleolus, **IRF** - intranucleolar replication focus. Scale bar (A-E) – 50 μm, (F-I) - 2 μm.

### Lowicryl sections are compatible with 2D and 3D single-molecule localization microscopy

We reasoned that the embedding of samples into Lowicryl and sectioning would reduce sample complexity (e.g. minimal thickness, sample immobilization) and improve SMLM characteristics. Using the Click-iT labelling protocol in Lowicryl sections, we obtained good photo-switching properties, averaging 128 780 localizations (n = 28 images, s.d. = 61 275; with the number of localizations normalized for the number of cells in the field of view) with minimal out-of-focus detections given the physical size of the specimen (500 nm). Besides localization microscopy metrics such as localization precision or the average number of localizations, we used Nano-J SQUIRREL to evaluate possible errors in image reconstruction (Fig. S1). Overall, we observed a high correlation between widefield and super-resolution images, with global resolution-scaled Pearson correlation coefficients for analysed images (RSP) being 0.96, 0.91 and 0.76 (Fig. S1 C1 - C3). The image resolution, calculated from the Fourier ring correlation analysis, was measured to be approximately 18 nm (n = 10; SD = 8.8 nm; Fig. S1 E1-E2), within the expected range (10 – 40 nm).

We hypothesized that the improved imaging characteristics for replication labelling on sections could be due to both reduction of sample complexity as well as good accessibility of the Alexa Fluor 647-azide conjugate to the epitopes embedded in the resin. To visualize the depth of penetration for the Alexa Fluor 647-azide conjugate into the resin we prepared and imaged 1000 nm (Fig. 4 A, B) as well as 500 nm (Fig. 4 C, D) thick sections of the root tissue. As seen in Fig. 4, we detected localizations from the entire volume of the section, visible in the XZ projections (Fig. 4 A, C) and 3D projections (Fig. 4 B, D). The number of localizations for 3D reconstructions is considerably lower than for 2D (35717 average localizations, n = 9 images, s.d. = 15234, as opposed to 128 780 average localizations in 2D), probably due to the filtering of a subset of localizations due to aberrant PSF deformation.

**Figure 4.**
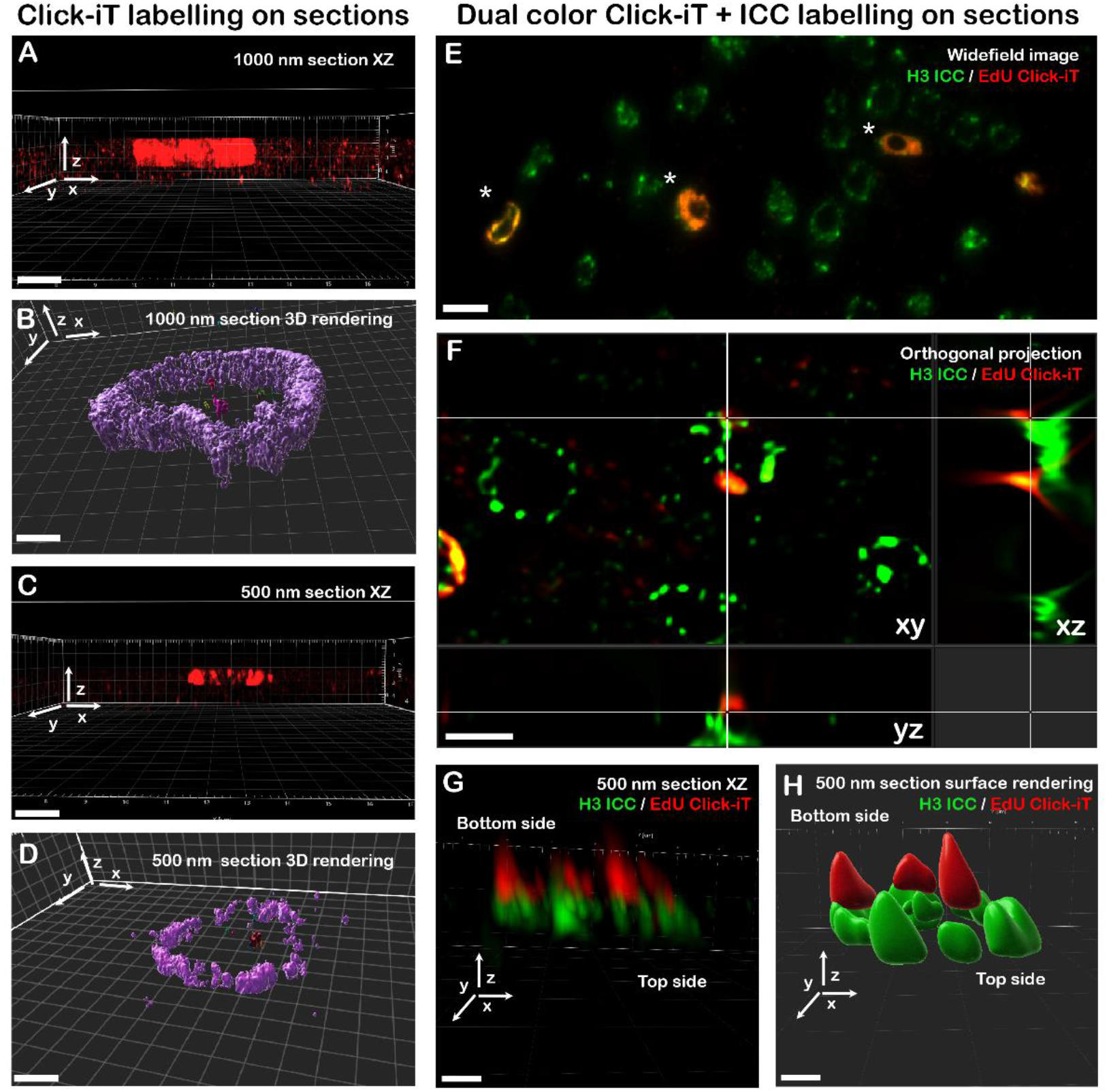
3D single-molecule localization microscopy and resin permeability on Lowicryl sections. Comparison of 1000 nm (**A, B**) and 500 nm (**C, D**) section labelling with Click-iT chemistry shown in the XZ projection (**A, C**) and 3D rendering (**B, D**). Dual labelling of histone H3 and DNA replication by EdU on 500 nm sections **E**), asterisks (*) highlight replicating cells labelled with EdU-AlexaFluor 647, widefield microscopy. Different depth penetration of the antibody and Click-iT labelling are shown in the orthogonal projection (**F**), with detailed views in **G** and surface rendering shown in **H**. Bottom side refers to the side of the section attached to the coverslip, the top side is exposed to buffer. Legend: **ICC** - immunocytochemistry. Scale bar: (**A**-**D**) – 1 μm (**E**) - 5 μm, (**F**) - 2 μm, (**G**-**H**) - 1 μm.

### Immunolabelling of histone H3 and phalloidin tagging of actin filaments on Lowicryl sections

After confirming that EdU-labelling coupled with Click-iT chemistry is efficient for tagging DNA (Fig. 2, Fig. 3A), and performs well in SMLM (Fig. 4 A-D, Fig. S1), we tested whether other labelling strategies can be used for Lowicryl sections. We found that dual labelling of chromatin with histone H3 by indirect immunofluorescence and DNA replication sites by Click-iT chemistry also yields sufficient levels of signal when applied on sections and can be used to demonstrate the proportion of EdU-tagged replicating cells in the cell population (Fig. 4E). Notably, labelling of chromatin with an anti-H3 antibody was restricted to the section surface (Fig. 4 F-H) with poor performance in SMLM in comparison to Click-iT labelling in terms of signal-to-noise ratio and the number of localizations (Fig. S2 A, B). The discrepancy in performance between H3 and EdU detection was likely due to several factors, including labelling density, fluorophore properties and limits in resin permeability for the relatively large IgG antibody (14 nm). We next probed the labelling of cellular structures with the low MW compound phalloidin conjugated to Alexa Fluor 647, which labels actin filaments in root sections. As seen in Figure S3, we detected strong labelling adjacent to cell walls with filaments extending into the cytoplasm (Fig. S3 - Insets), suggesting that detecting cellular structures with low MW compounds is feasible for Lowicryl resin-embedded tissue sections. Overall, we have demonstrated that Lowicryl embedding of samples is compatible with SMLM and allows for efficient labelling with Click-iT chemistry, low MW compounds or immunolabelling. Next, we used the established protocol to study the links between intranucleolar replication and changes in nucleolar architecture.

### Alterations in the nucleolar architecture and intranucleolar replication foci size in the *fas1* and *nuc1 Arabidopsis thaliana* mutants

Replication labelling is unique in allowing the discrimination of the S-phase progression, altogether with the identification of intranucleolar replication foci (IRFs). In combination with TEM, we were able to link the distribution of IRFs with the architecture of the nucleolus, especially the presence and size of fibrillar centers (FCs), not visible in light fluorescence microscopy. From pooled data from all the experiments, we observed that cells in the late-S-phase did not show IRFs (Fig. 5 A, H) even though they did display FCs (88,8%; n = 9). For cells in the early-S-phase (Fig. 5 B, C), we mostly found nuclei without prominent IRFs or FCs or containing both structures (Fig. 5C, D; 54% and 32%, respectively, n = 28). The quantification revealed (Fig. 5G, H) that the presence of FCs together with IRFs is specific to early-S-phase progression. The co-occurrence of FCs and IRFs, altogether with partial colocalization (Fig. 5C1 - 5C2) suggests that fibrillar centers are the sites of rDNA replication.

**Figure 5.**
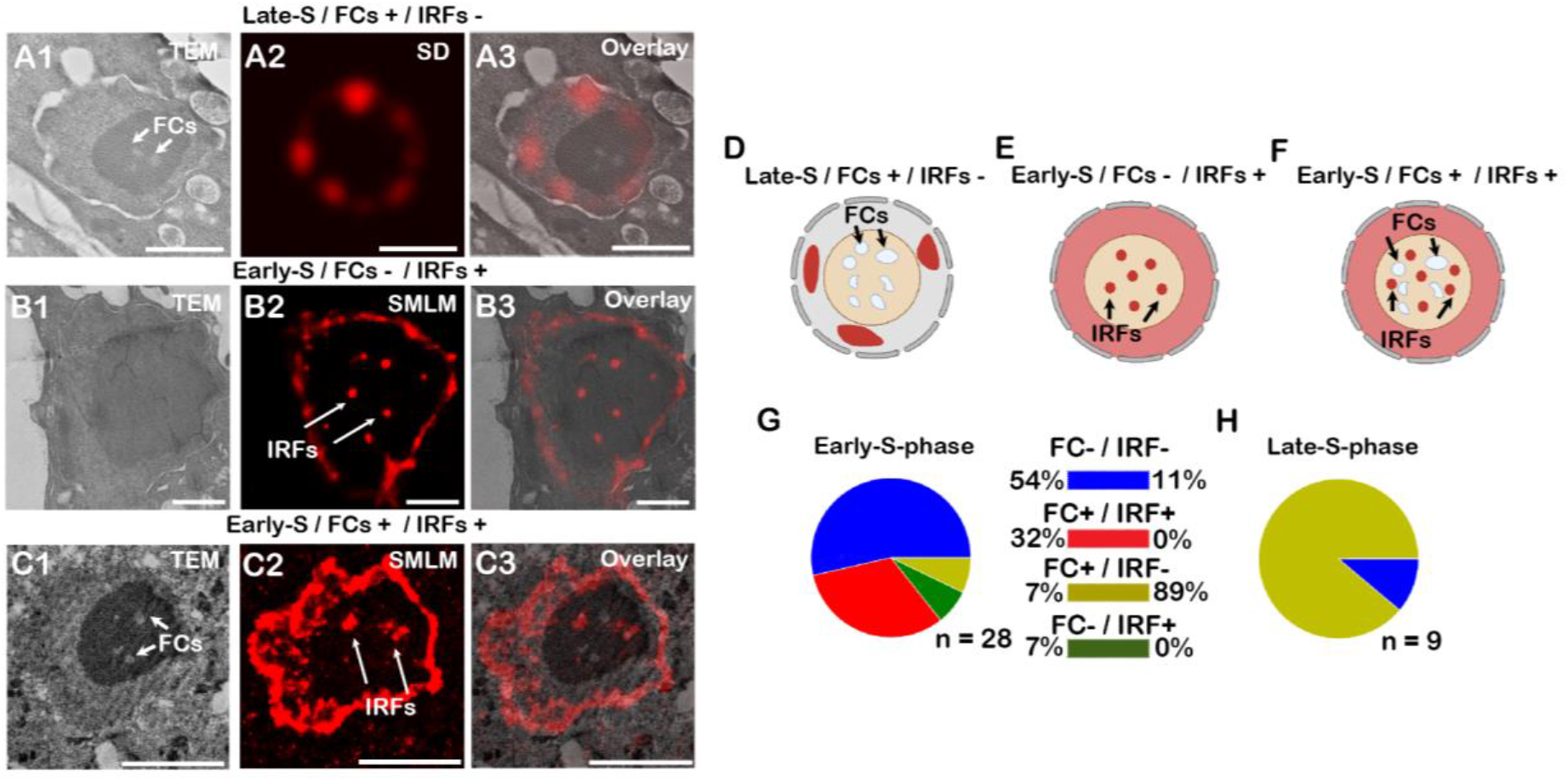
Analysis of the correlation between S-phase progression and nucleolar architecture. **A1 - A3)** CLEM image of a nucleus in the late-S-phase (**A1** - TEM, **A2** - SD, **A3**-overlay image), with a schematic depiction in **D**. Examples of early to mid-S phase replicating nuclei (**B1**-**B3**; **C1**-**C3**, schematically in **E, F**), without and with visible FCs (**B** and **C**, respectively). Quantification of the replicating profiles (**G** - early and **H** - late) in relation to the presence of FCs and IRFs, n refers to the number of cells analysed, pooled dataset from both wild-type and mutant plants. Legend: **SD** - spinning disk, **TEM** - transmission electron microscopy, **FC** - fibrillar centers, **IRFs** - intranucleolar replication foci, **SMLM** - single-molecule localization microscopy. Scale bar (A-C) - 2 μm.

To test whether the size of IRFs and FCs differs between wt and *fas1* or *nuc1* mutant plants, we calculated the average diameter of IRFs from SMLM reconstructions (Fig. 6A) and FCs from TEM data (Fig. 6C). Mapping of IRFs from SMLM reconstructions revealed the average diameter of IRFs to be 191.5 (n = 29; s.d. = 53.1), 279.3 (n = 22; s.d. = 78.2) and 268.8 (n = 17; s.d. = 58.9) nm for wt, *fas1* and *nuc1* plants, respectively (Fig. 6B). For FCs, we found an average diameter of 332.5 (n = 38; s.d. = 66.1), 244.1 (n = 39; s.d. = 62.9) and 268.8 nm (n = 17; s.d. = 52.8) in the wt, *fas1* and *nuc*1 plants, respectively (Fig. 6D). The average size of IRFs in *fas1* and *nuc1* mutants was significantly higher relative to IRFs in the wt (Kruskal-Wallis H-test; p = 1.01 × 10^−5^, H-statistic = 23; Wilcoxon rank sum test adjusted p-value = 5.1 × 10^−5^ between wt and *fas1*; and p = 7.9 × 10^−5^ between wt and *nuc1*). Likewise, mutant plants had a reduced size of FCs (Kruskal-Wallis H-test; p = 4.2 × 10^−7^; H-statistic = 29.4; Wilcoxon rank sum test adjusted p-value = 1.1 × 10^−6^ between wt and *fas1*; p = 5.3 × 10^−5^ between wt and *nuc1*).

**Figure 6.**
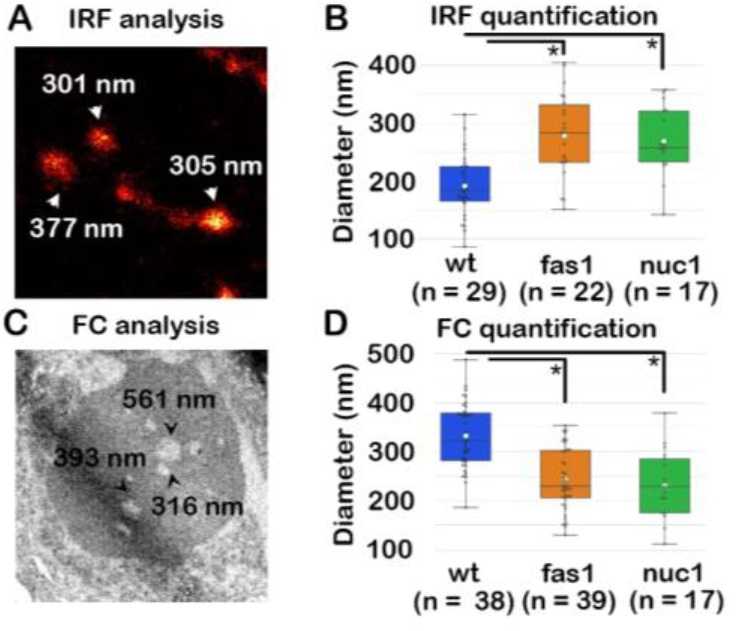
Quantification of the FC and IRF diameter in wild-type and mutant *A. thaliana*. Quantification of the size of IRFs (**A, B**) in SMLM micrographs. Quantification of FCs from TEM micrographs in (**C, D**). For IRF and FC quantification (**B, D**), n refers to the number of structures analyzed. Statistical significance was evaluated based on the non-parametric Kruskal-Wallis H-test. Legend: **FC** - fibrillar centers, **IRFs** - intranucleolar replication foci.

Besides changes in the size of FCs and IRFs, *nuc 1* mutants showed altered nucleolar morphology, in line with previous reports (Pontvianne et al., 2007). The phenotypic changes such as less pronounced FCs with an atypical shape and weaker contrast in the nucleoli occur irrespective of fixation (Fig. S4A, S4C) and embedding (Fig. S4B, S4D) conditions, since we detect them in both Spurr’s and Lowicryl embedded sections. Altogether, we note important differences in the replication of nucleolar material, parallel with changes in the nucleolar structure in mutant plants.

### Analysis of replication progression in wild-type, *fas1* and *nuc1* mutant plants using SMLM imaging

Replication of the nuclear genetic material is a multi-step process that involves chromatin remodeling that is required for access to the replisome complexes by the DNA polymerase. To investigate whether the possible changes in replication progression in histone chaperone mutants such as *fas1* or *nuc1* manifest in the nuclear replication pattern, we conducted density-based clustering analysis of SMLM data acquired on semi-thin sections. First, we performed SMLM image reconstruction in SMAP (Fig. 7A), then filtered the localizations by clustering density (Fig. 7B) and performed density-based spatial clustering of applications with noise (DBSCAN) (Fig. 7C). We chose DBSCAN clustering, as it a robust technique that does not require specification of the number of clusters, and is not sensitive to cluster shape (Wu et al., 2020). In all conditions, we observed a spectrum of replication patterns, ranging from dozens up to hundreds of clusters per nucleus observed after thresholding. We did not observe statistically significant (Kruskal-Wallis p-value = 0.078) differences between replication foci clustering in wt, *fas*1 or *nuc*1 plants (Fig. 7D).

**Figure 7.**
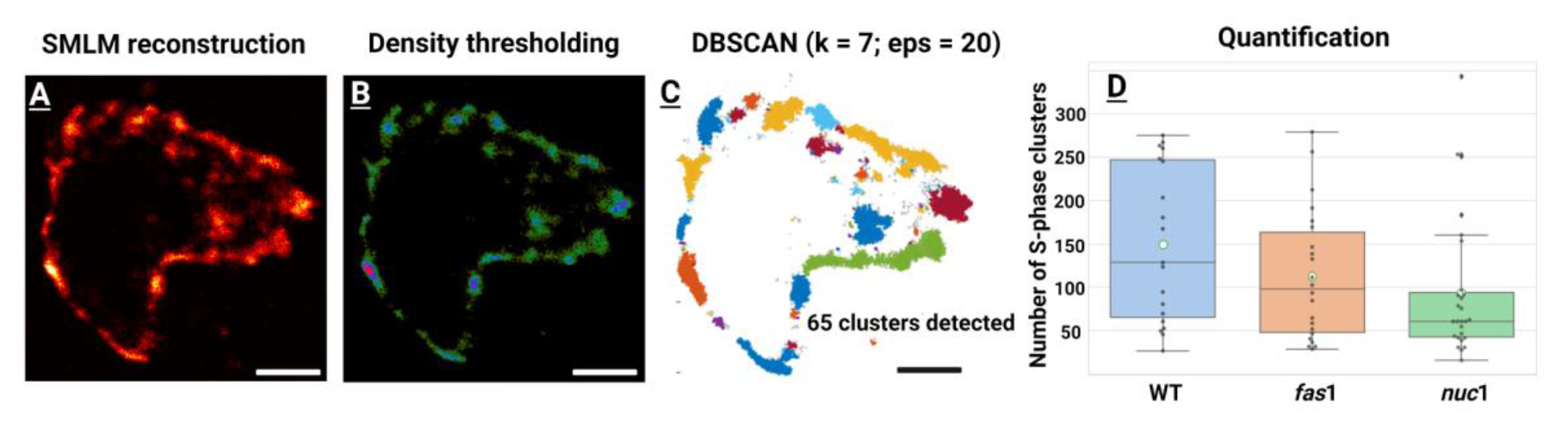
Clustering-based analysis of replication progression on SMLM data. Super-resolution images were reconstructed in the SMAP software (**A**). Image thresholding using cluster density calculator (**B**) to eliminate localizations from low-density regions. **C**) Detection of clusters using the DBSCAN algorithm. Quantifying the number of clusters in wt, *fas*1 and *nuc*1 mutants using the Kruskal-Wallis H-test (**D**). Legend: **k** - minimum objects in the neighbourhood, **eps** - neighbourhood radius. Scale bar - 1 μm.

## Discussion

Microscopy has always represented a fundamental tool for the in-depth analysis of internal cellular structures. Unfortunately, the application of advanced imaging techniques in plants (e.g. SMLM or CLEM approaches) often lags behind the implementations in other model systems. This is due mostly to challenges in addressing the complexity and composition of plant tissues, often requiring customization of protocols and imaginative ways to implement otherwise common labelling methods. The applications of SMLM in plants are rare (Dong et al., 2015; Mass et al., 2020; Kubalová et al., 2021; Schnorrenberg et al., 2020), with groups often opting for less demanding high-end techniques such as SIM or 3D-SIM (reviewed in Schubert, 2017). In our case, we first tested imaging plant roots using SMLM in a whole-mount setup, with sub-optimal performance. Without immobilization on the coverslip, the roots tended to detach from the coverslip and move when submerged in the buffer, with poor-photoswitching properties during acquisition. We also observed significant Alexa Fluor 647 photobleaching (the average number of localizations = 22311; n = 6; s.d. = 9058 over 10000 time frame acquisition; data not shown). Correlative techniques are generally faced with similar challenges in sample preparation and requirements for protocol optimization. While it has been shown previously that alternate sectioning of embedded plant tissues for fluorescence and electron microscopy can be used for correlative analysis (Bell et al., 2009), the implementation of super-resolution microscopy in tandem with electron microscopy in plants, especially in conjunction with the unique properties provided by bio-orthogonal chemistry labelling has not been shown previously.

Bridging electron microscopy with super-resolution analysis of cellular ultrastructure on sections was the key technical focus of this study. We present a workflow for the correlative imaging of plant tissues that combines chemical fixation, Lowicryl embedding, on-section labelling for electron microscopy and super-resolution microscopy, focused on studying DNA replication and nucleolar organization in cells. Our workflow has several advantages compared to approaches that use isolated nuclei and conventional labelling methods, both in terms of biological relevance and technical / imaging capabilities. First of all, imaging of plant sections enables us to estimate cell identity based on the position of the cell within the tissue. In our case, we focused primarily on the nucleolar architecture and replication profiles in cells of the epidermal layers from the meristematic and elongation zones. Second, the sectioning of tissues, coupled with the specific Click-iT labelling results in a lower fluorescence background which is critical for SMLM imaging, as it translates directly into image reconstruction quality and improved resolution.

In the context of replication labelling presented in this study, EdU incorporates selectively into replicating DNA and the subsequent Click-iT reaction mediates the covalent binding of the conjugated fluorophore to ethynyl (for an overview of Click-chemistry see Sletten & Bertozzi, 2009 and Best, 2009). The incorporation of bio-orthogonal (Click-iT) labelling into the workflow is important with respect to the fixation conditions and SMLM performance. The fixation of samples with glutaraldehyde is necessary for the preservation of the fine ultrastructure of subcellular features, though it can result in structural alterations, masking of some immune-epitopes and cause background fluorescence. We show that the Click-iT reaction between the azide and ethynyl moieties is not hindered by fixation with up to 3% glutaraldehyde. Moreover, the Click-iT reaction is not prevented by either post-contrasting with osmium tetroxide or embedding into Spurr’s resin, although the high background on osmium tetroxide post-contrasted samples interferes with SMLM imaging. This suggests that Lowicryl embedding is the optimal choice for applications involving advanced microscopy techniques. In terms of SMLM, Click-iT labelling introduces virtually no linkage error (the distance between the fluorophore tag from the actual position of the epitope), which makes it an optimal strategy for tagging structures. Importantly, Click-iT chemistry is not restricted to labelling nucleic acids and can be successfully applied to tag proteins of interest through the introduction of non-canonical amino acids (Nikic et al., 2015; Parker and Pratt, 2020). In recent publications, Click-iT chemistry has been introduced into CLEM workflows and applied to tracking intracellular trafficking or lipids in bacteria (Adrian et al., 2021; Peters et al., 2021), showing the promise of this approach in imaging applications.

To demonstrate the utility of the established correlative workflow presented in this work, we applied it to examine the nucleolar ultrastructure during replication. Using a CLEM approach in this study was essential to determine whether the presence of FCs correlates with the detection of IRFs in the early S phase. We show that the replication of intranucleolar DNA is restricted to the early phases of S-phase progression, sometimes concomitant with the detection of fibrillar centers. We acknowledge that the distinction between early S-phase and gradual progression towards mid S-phase is not always easy to evaluate from microscopy data without flow cytometry, though the absence of IRFs in late S-phase cells is evident. Moreover, our results show statistically significant changes in the nucleolar ultrastructure in the *fas1* and *nuc1* plants, as reflected in the decreased size of fibrillar centers (FCs) relative to the wild type. While we cannot assume this decrease in size is the causal factor for the increase of the size of intranucleolar replication foci (IRFs), it is clear that the changes in nucleolar ultrastructure impact physiological processes in the nucleolus, notably replication as detected through DNA replication labelling. We hypothesize that the increase in the size of IRFs is the result of the relaxed chromatin configuration, which might be related to the altered methylation status of rDNA and its decompaction (*nuc*1 mutants; Pontvianne et al., 2007), different levels of histone variants present in rDNA and/or general redistribution of rDNA genes in the nucleolus (Kolarova et al., 2020; Kutashev et al., 2021). Our analysis of replication foci clustering in the nucleus did not show significant differences between the wild-type and *fas*1 or *nuc*1 mutant plants, in line with previous observations suggesting that the S-phase progression is not delayed in the *fas*1 background (Eekhout et al., 2021).

While we believe that the experimental approach presented in this paper is an important advance in correlative and super-resolution imaging of plant nuclei, it has certain limitations. First of all, while we can analyse the same cells in both TEM and SR imaging, there is always an offset in the axial plane, since we are imaging two different adjacent sections (500 nm section for SMLM and 70 nm section for TEM). When we tested Click-IT labelling on TEM grid immobilized 70 nm sections, the fluorescence signal was too low for confocal or super-resolution imaging (data not shown). The second important limitation is tied to labelling. We make the case that Lowicryl sections are permeable for low molecular weight fluorophore conjugates, but detecting proteins with classical antibodies is limited since they cannot penetrate Lowicryl (Stierhof and Schwarz, 1989, Fig. 4 F-H). However, this limitation may be overcome by the use of SNAP-tagged or HALO-tagged proteins expressed in plants (Iwatate et al., 2020) and their subsequent detection in sections using fluorescently labelled SNAP or HALO ligands characteristic for their low MW and efficient penetration. Lastly, the correlation between SMLM and EM images is not trivial, due to sample deformation and stretching during processing. Some shrinking occurs during water removal from the hydrated tissue – the processing step necessary for tissue infiltration with resin. Partial shrinking of sections also occurs during their cutting in the ultramicrotome (most apparent along the cutting axis) while stretching of sections may occur during section mounting on supporting EM grids or glass coverslips and during exposure to a high dose of focused electron beam inside the TEM. Partial mismatch of the overlay of the light/fluorescence image (500 nm section) and the corresponding TEM image (70 nm section) must also be expected due to fact that both are 2D-projections of a (partial) 3D-volume of a different thickness and the ultrathin TEM section is an adjacent section (continuum) of the semithin section.

Overall, we show that the replication labelling in plant cells is feasible on Lowicryl sections with optimal properties for single-molecule localization microscopy, allowing the analysis of DNA replication in a quantitative manner with the ultrastructure of the nucleolus provided through correlation with TEM imaging. We envision findings presented in this paper open the path to novel applications of resin embedding in tandem with super-resolution microscopy, including relevant plant crop models. Based on our findings, we plan to test the utility of the workflow for other applications, including combinations with fluorescence in-situ hybridization for the detection of specific genomic segments (e.g. telomeric repeats), complemented with chromatin contrasting based on local 3 ‘3-di-amino-benzidine (DAB) polymerization (Ou et al., 2017).

## Acknowledgments

We acknowledge the core facility CELLIM supported by the Czech-BioImaging large RI project (LM2023050 funded by MEYS CR) for its support in obtaining scientific data presented in this paper and the BIOCEV imaging facility for their support. We acknowledge Imaging Methods Core Facility at BIOCEV, supported by the MEYS CR (LM2023050 Czech-BioImaging) and for their support & assistance in this work. The project was funded by the European Regional Development Fund – Project ‘SINGING PLANT’ CZ.02.1.01/0.0/0.0/16_026/0008446. We acknowledge the Czech Bioimaging initiative for providing access and financial support for advanced imaging techniques. Figures 1 and 5 have been co-created with BioRender.com

## Author contributions

MF and LK performed all experiments and sample preparation. MF, LK, JM, JP and EM performed electron and super-resolution microscopy. MF conducted data analysis. MF, LK, DL, ME, MD and JF performed experimental design, manuscript preparation and provided technical background and resources for the study.

## Supplementary information

**Figure S1.**
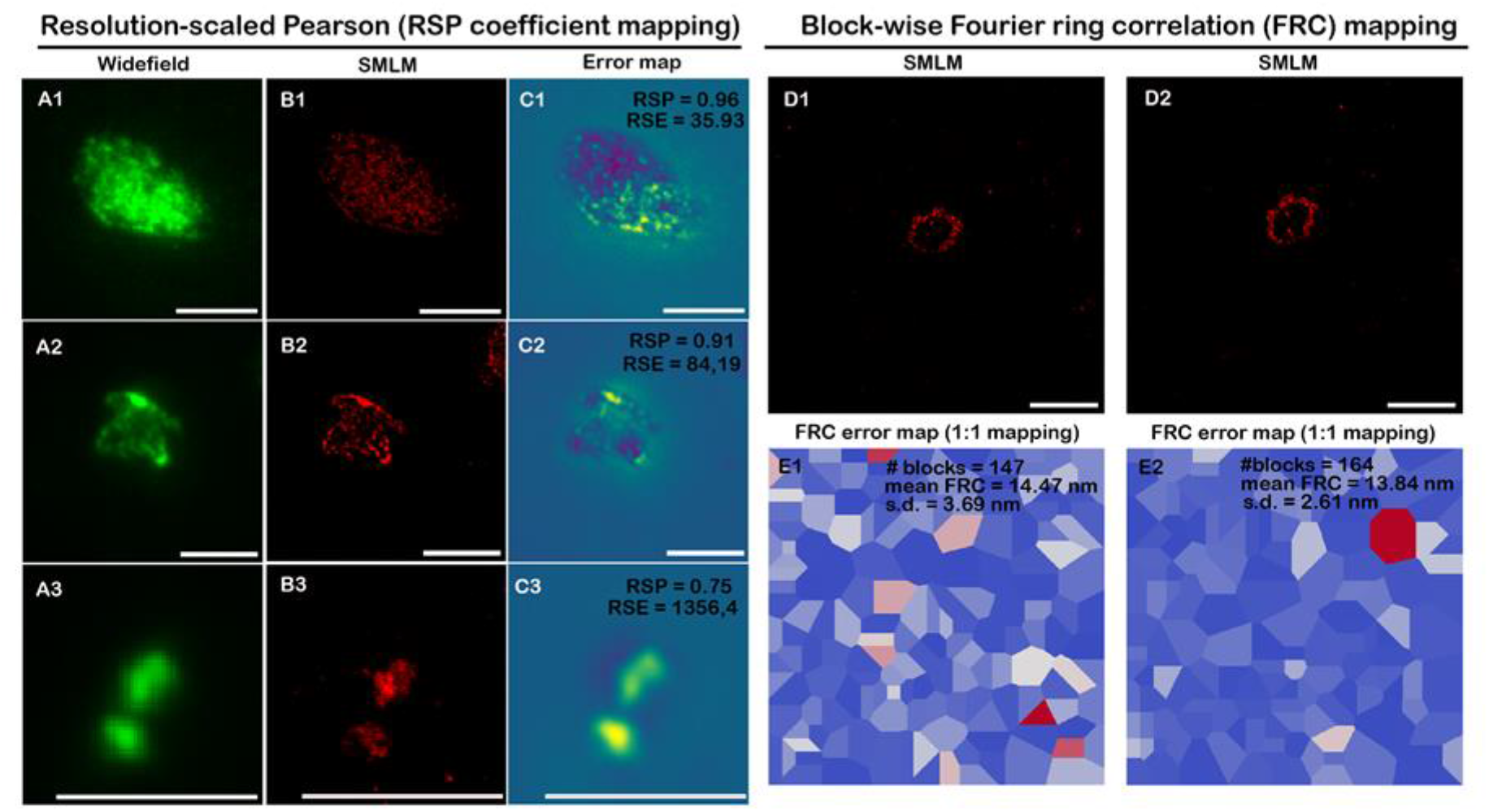
Quantitative error mapping of SMLM reconstructions from sections in Lowicryl. Analysis of reconstruction errors using NanoJ-SQUIRREL. Global resolution-scaled Pearson coefficient (RSP) is calculated from the widefield (**A**) and reconstructed SMLM (**B**) images and provides an image quality metric (values ranging from -1 to 1, indicating full anti-correlation and correlation, respectively - **C1, C2, C3**). Splitting of the original time series used for reconstruction (reconstruction in **D1, D2**) into different substacks can be used to calculate the Fourier ring correlation, used to estimate image resolution globally (mean FRC values), with local heterogeneities visible from the error maps (**E1, E2**). Scale bar - 5 μm.

**Figure S2.**
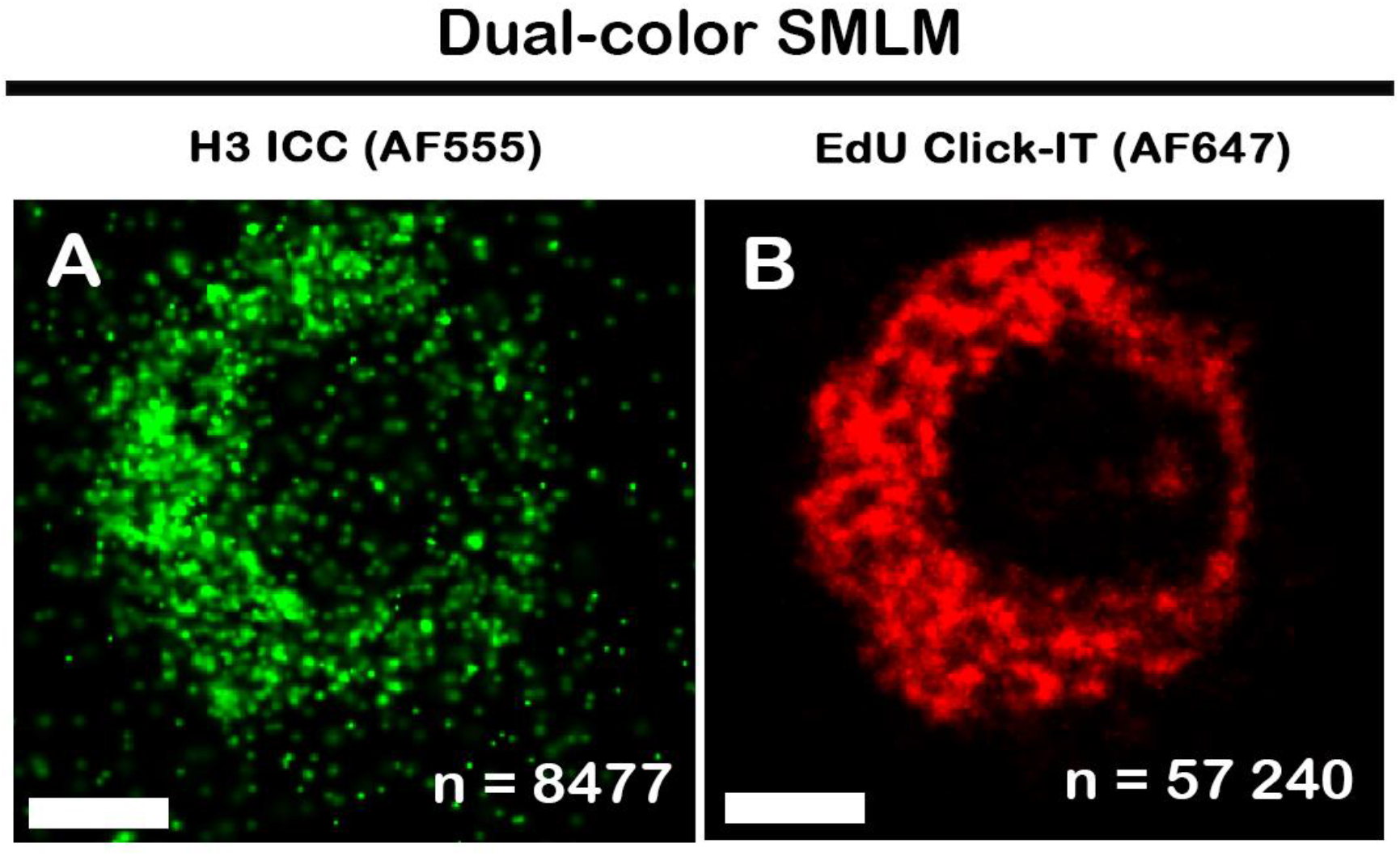
Dual-color super-resolution microscopy on sections. Antibody labelling of the H3 histone, with Alexa Fluor 555 secondary antibody (**A**). Click-iT labelling with EdU and Alexa Fluor 647 detection in (**B**). The number of localizations is indicated at the bottom left of each image, after filtering and image reconstruction in the ZEN Black software. **ICC** - immunocytochemistry. Scale bar - 1 μm.

**Figure S3.**
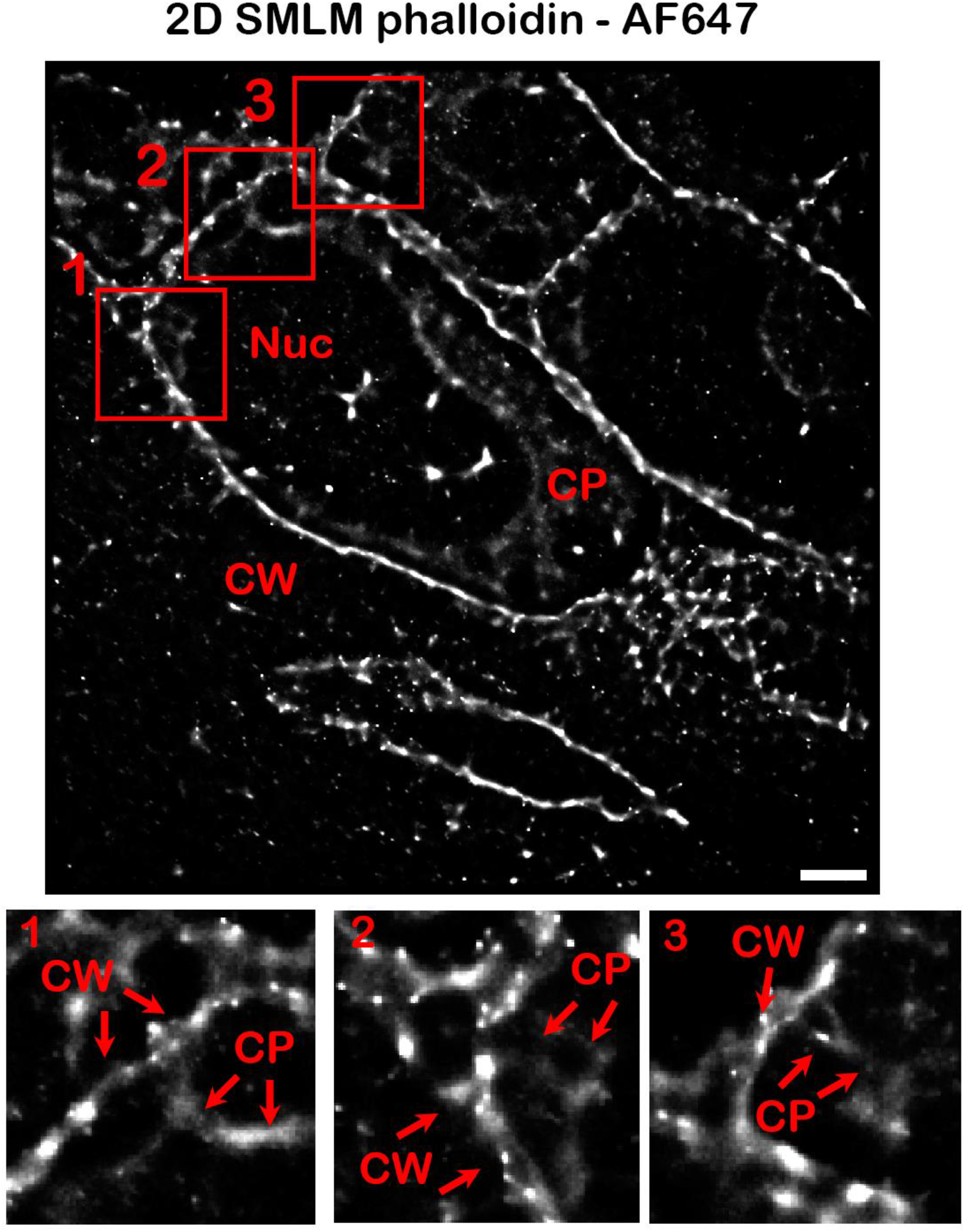
Detection of actin ultrastructure in Lowicryl sections. SMLM analysis of actin in root tip cells (**A**). Insets (**1**-**3**) display actin branching from the cell wall fraction (**CW**) to the cytoplasmic structures (**CP**). **CW** - cell wall fraction, **CP** - cytoplasmic fraction. Scale bar - 2 μm.

**Figure S4.**
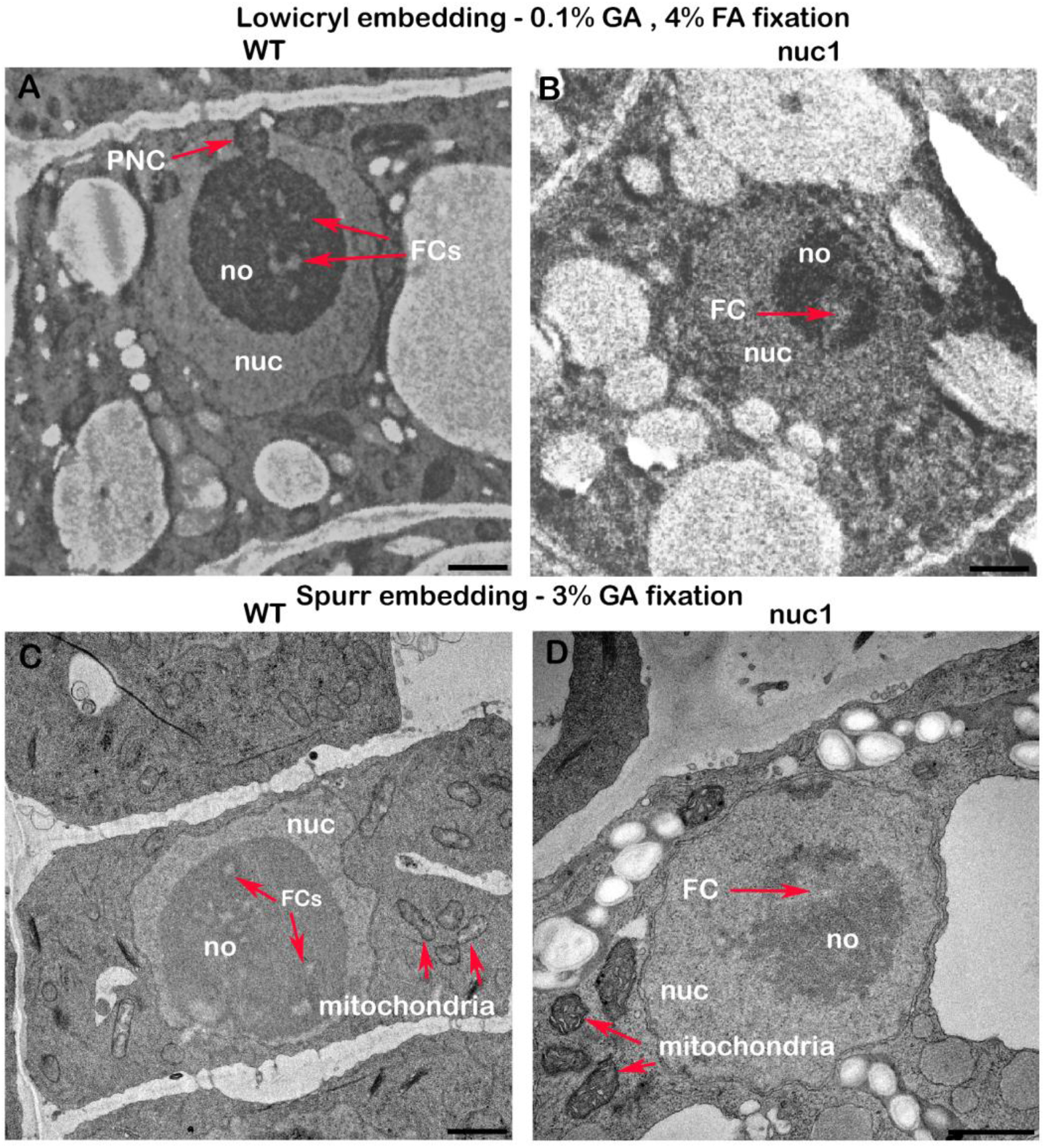
Nucleolar ultrastructure in wild-type and *nucleolin 1* mutants. Differences in the nucleolar ultrastructure between wild type (**A, C**) and *nuc1* (**B, D**) mutants in Lowicryl-embedded (**A, B**) and Spurr’s-embedded (**C, D**) samples. Epidermal cells of wild-type plants show typical features of nucleoli, including perinucleolar clusters and clearly delineated fibrillar centers (**A**). Nucleoli of *nucleolin 1* mutants show weak contrast in fibrillar centers with indistinct, diffuse edges (**B**). These features can be observed also in samples after stronger fixation conditions (3% GA) and Spurr’s embedding (**C, D**). **Nuc** - nucleus, **no** - nucleolus, **PNC** - perinucleolar cluster, **FC** - fibrillar center. Scale bar - 1 μm.

