## Supplementary files for "In-section Click-iT detection and super-resolution CLEM: Shedding light on nucleolar ultrastructure and S-phase progression in plants"

### Supplementary information

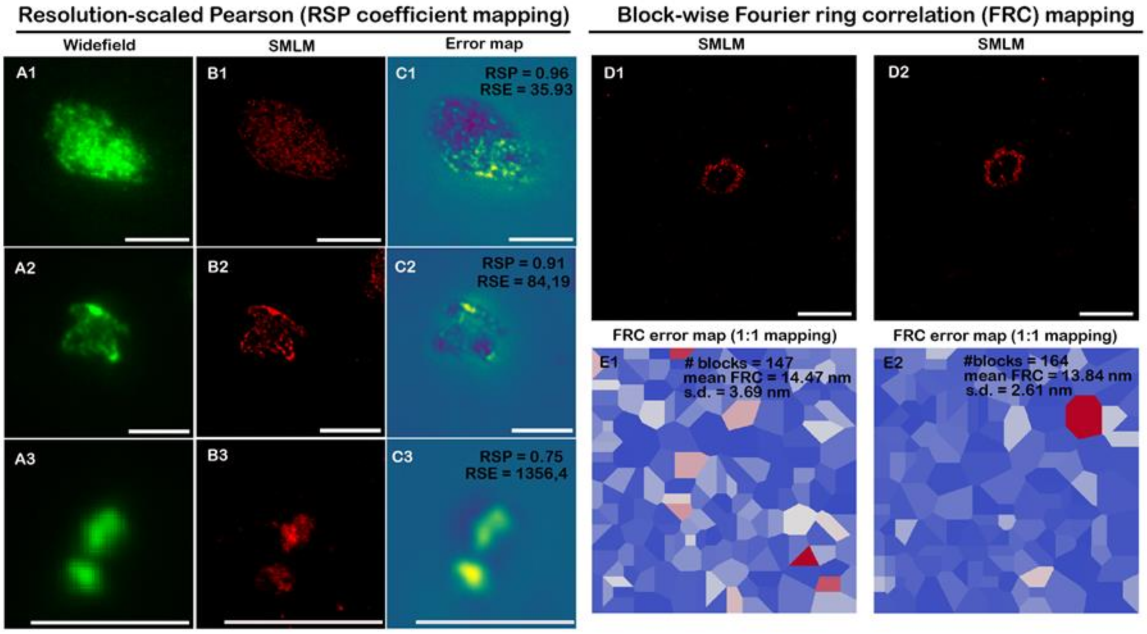


**<u>Figure S1. Quantitative error mapping of SMLM reconstructions from sections in Lowicryl</u>**

Analysis of reconstruction errors using NanoJ-SQUIRREL. Global resolution-scaled Pearson coefficient (RSP) is calculated from the widefield (**A**) and reconstructed SMLM (**B**) images and provides an image quality metric (values ranging from -1 to 1, indicating full anti-correlation and correlation, respectively - **C1**, **C2**, **C3**). Splitting of the original time series used for reconstruction (reconstruction in **D1**, **D2**) into different substacks can be used to calculate the Fourier ring correlation, used to estimate image resolution globally (mean FRC values), with local heterogeneities visible from the error maps (**E1**, **E2**). Scale bar - 5 µm.

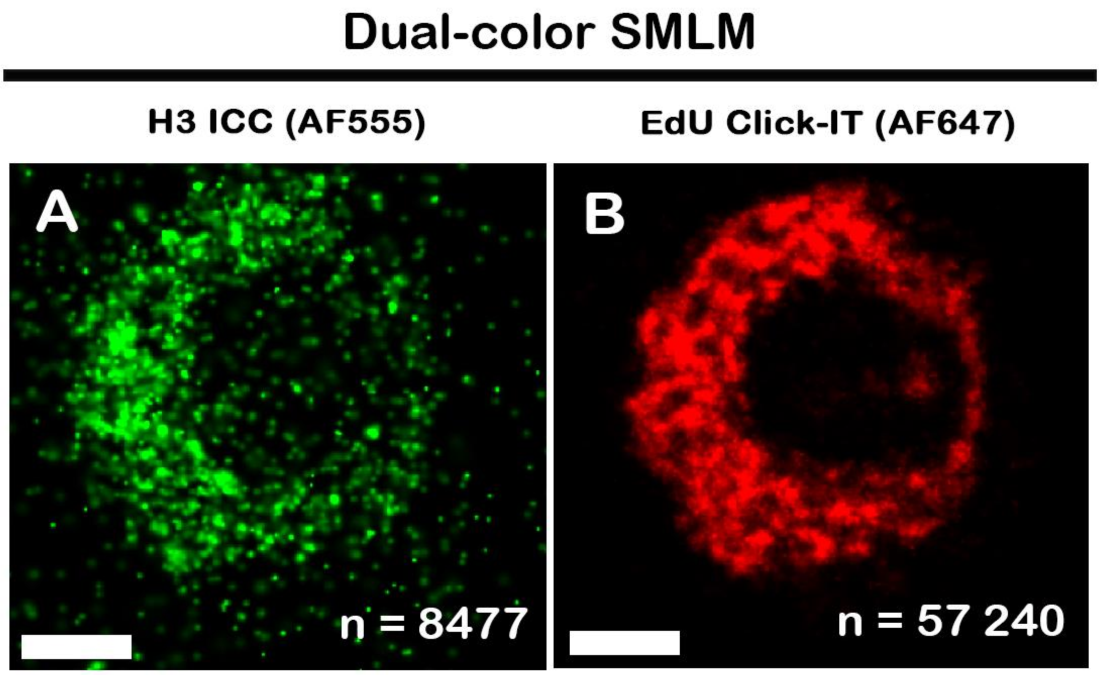


**Figure S2. Dual-color super-resolution microscopy on sections.**

Antibody labelling of the H3 histone, with Alexa Fluor 555 secondary antibody (**A**). Click-iT labelling with EdU and Alexa Fluor 647 detection in (**B**). The number of localizations is indicated at the bottom left of each image, after filtering and image reconstruction in the ZEN Black software. **ICC** - immunocytochemistry. Scale bar - 1 μm.

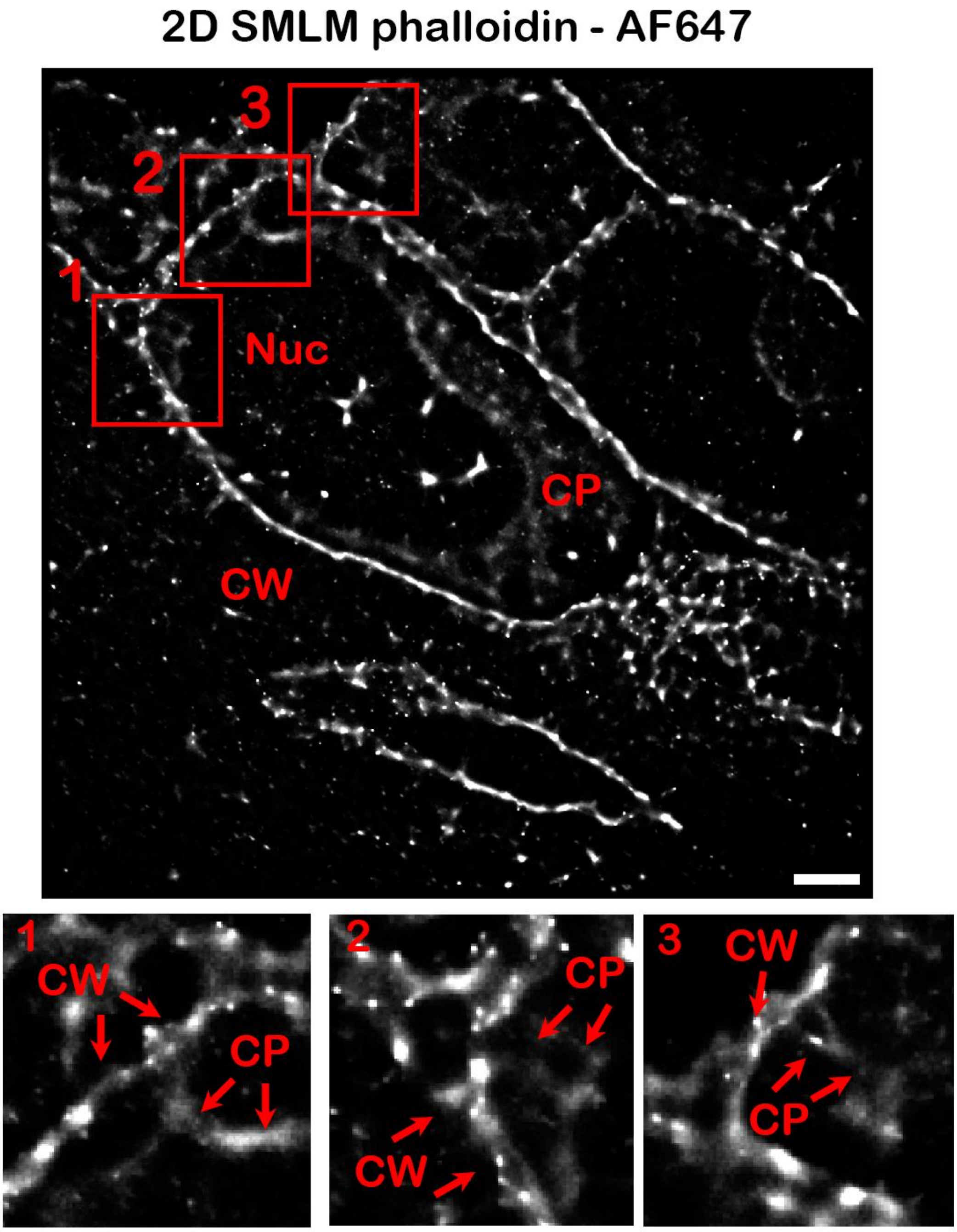


**<u>Figure S3. Detection of actin ultrastructure in Lowicryl sections</u>**

SMLM analysis of actin in root tip cells (**A**). Insets (**1**-**3**) display actin branching from the cell wall fraction (**CW**) to the cytoplasmic structures (**CP**). **CW** - cell wall fraction, **CP** - cytoplasmic fraction. Scale bar - 2 µm.

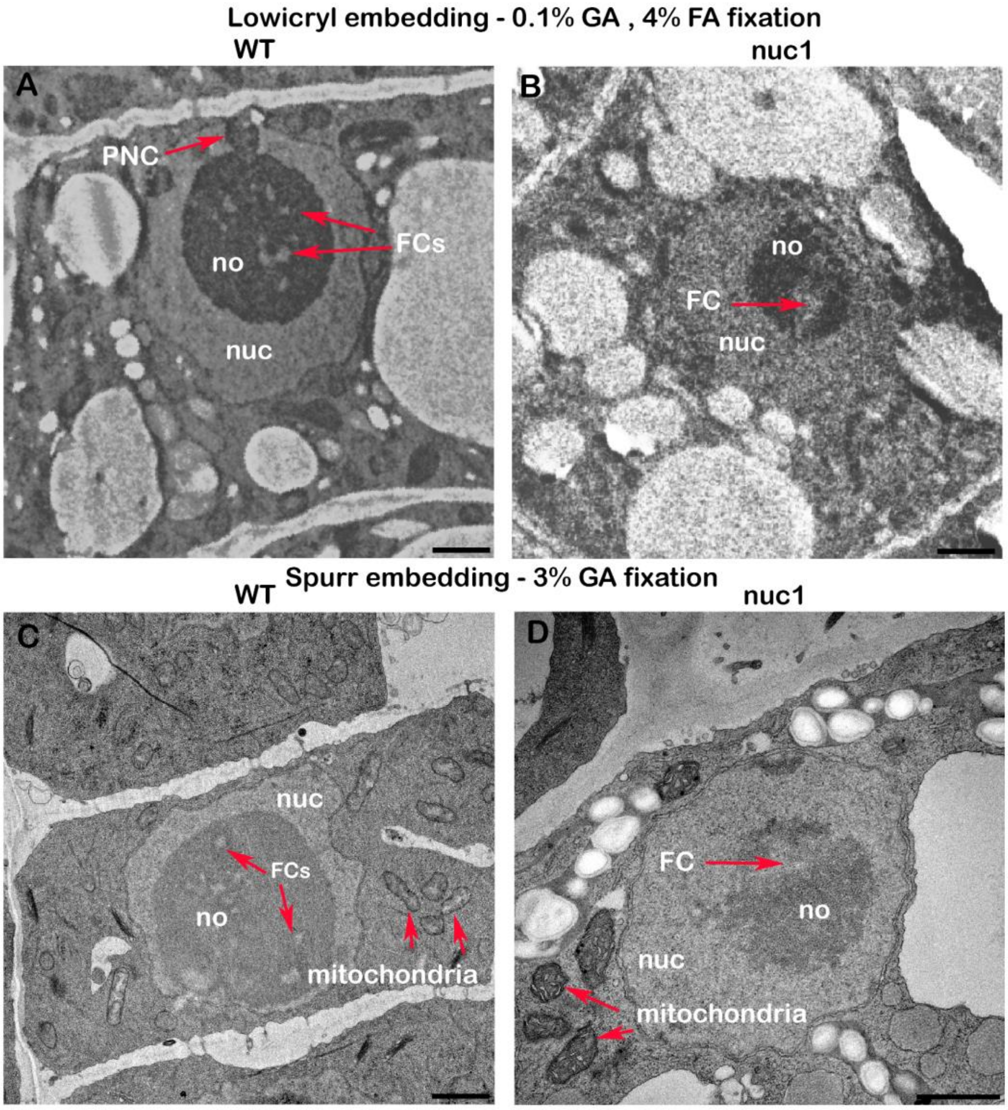


### Figure S4. Nucleolar ultrastructure in wild-type and *nucleolin 1* mutants

Differences in the nucleolar ultrastructure between wild type (**A**, **C**) and *nuc1* (**B**, **D**) mutants in Lowicryl-embedded (**A**, **B**) and Spurr's-embedded (**C**, **D**) samples. Epidermal cells of wild-type plants show typical features of nucleoli, including perinucleolar clusters and clearly delineated fibrillar centers (**A**). Nucleoli of *nucleolin 1* mutants show weak contrast in fibrillar centers with indistinct, diffuse edges (**B**). These features can be observed also in samples after stronger fixation conditions (3% GA) and Spurr's embedding (**C**, **D**). **Nuc** - nucleus, **no** - nucleolus, **PNC** - perinucleolar cluster, **FC** - fibrillar center. Scale bar - 1 µm.
